# Slivka and Slivka-bio: a lightweight framework for presenting executables as web services and its application in bioinformatics

**DOI:** 10.64898/2026.05.23.727389

**Authors:** Mateusz Warowny, Thomas Down, Stuart A. MacGowan, Kiran Mukhyala, Geoffrey J. Barton, James B. Procter

**Affiliations:** Faculty of Life Sciences, University of Dundee, Dow Street, Dundee, DD1 5EH, Scotland, UK; Department of Structural Biology, Genentech, 1 DNA Way, South San Francisco, CA 94080, USA

## Abstract

**Motivation:** Execution of code is critical for computational biology, but technical requirements can prevent others from running it. Public web-apps and services thus remain the most effective way to make code accessible, but no fully reusable infrastructure exists to help researchers do this.

**Results:** We developed Slivka to enable easy provision of robust HTTP-based execution services backed by local or distributed hardware; accessible via curl and dedicated clients. We demonstrate it with Slivka-bio, which provides semantically annotated services for Jalview 2.12 (https://www.jalview.org/development/jalview_develop/) and includes 15+ tools for protein and RNA analysis. Slivka has been in production in academic and industry environments for 5 years and ran more than 1.5M jobs.

**Availability and Implementation:** Slivka and Slivka-bio are released under the Apache 2.0 License. Slivka-bio public instance at https://www.compbio.dundee.ac.uk/slivka with links to documentation, docker containers, and github repositories for Slivka-bio and Slivka.

## 1. Introduction

The rate at which new computational tools are developed is rapidly increasing but there can be a barrier to placing these tools in the hands of people who need them. Despite improvements in software packaging and availability of cloud-based computational resources, bespoke websites that allow a tool to be executed are still the simplest to ensure a tool becomes used and cited (Song & Kurgan, 2023). However, website maintenance can quickly become a burden, since they must remain operational as usage demands increase and technology changes; and Ősz et al. (2019) found that 15% of services become unavailable 3 years after publication. For tools that require large databases or proprietary resources, it is also important to provide programmatic access, so researchers can employ them as part of workflows. Web based Application Programming Interfaces (APIs) are central to the modern web, and many approaches exist for their creation. However, detailed technical knowledge is often required, and relatively few lightweight, re-usable systems exist that allow researchers to create a standards compliant API to execute their tool. To these ends, we describe Slivka – a system designed to make it easy to create web services for computational tools that are secure, straightforward to administer, and able to take advantage of existing research computing infrastructure.

We demonstrate Slivka’s capabilities with Slivka-bio, an easy to install Slivka service suite for bioinformatics. Slivka-bio services are described with EDAM(Ison et al., 2013; Black et al., 2022) annotations, which facilitates their automatic discovery with Jalview 2.12, an enhanced version of the Jalview (Waterhouse et al., 2009) interactive multiple sequence alignment, analysis, and visualisation platform. We also describe Czekolada – a graphical interface for Slivka servers that has allowed life scientists exploit to a range of computational tools for drug discovery which would otherwise be hard to access via command line.

### 1.1 Background

Web-based tools enable users to remotely execute code. They often also provide sophisticated ‘user-friendly’ user interfaces (web-UIs). At their core, these sites provide a way for users to input data, adjust parameters, submit and monitor progress, and download or view results.

Tool web sites that provide a web-based API also allow data upload, execution and results retrieval from scripts, enabling them to be employed as part of a larger workflow. Similarly, a web-based system might provide access to several tools. In fact, practically every research community in computational science has developed systems that allow remote tool execution (Andronico et al., 2011). In this work, this capability is referred to as ‘Execution as a Service’ (EaaS). Most recently, a number of systems have been developed to provide EaaS-like capabilities through ‘Model Context Protocol’ (MCP)(“Introducing computer use, a new Claude 3.5 Sonnet, and Claude 3.5 Haiku \ Anthropic”; “Model Context Protocol (MCP) Specification”). MCP is a JSON-RPC based API standard designed to expose tools and other capabilities to Large Language Model (LLM) and Agentic Artifical Intelligence (Agentic AI) systems. MCP essentially enables the same functionalities as other EaaS systems, and there are a number of reasons why future bioinformatics systems should support MCP(Flotho et al., 2026). However, MCP is a heavy-weight state-full protocol which has been optimised for AI clients, rather than simple programmatic access over HTTP. This means that any web-based user interface for an MCP server needs additional logic (most likely exposed as an intermediate API) to meet the additional expectations that MCP places on client systems (e.g. provision of data via a ‘virtual file system’). However, our goal is to provide a reusable EaaS API that does not require intermediate logic to serve both programmatic and UI based clients. For the majority of this paper we thus focus on the design of ‘traditional’ EaaS systems, and reserve further analysis of of EaaS *via* MCP to the Discussion (Section 5. Discussion).

In the Life sciences, Galaxy is one of the best known EaaS systems. It provides a sophisticated web user interface and web-API, accessible with dedicated clients such as Planemo(Bray et al., 2023). Very widely used EaaS systems are also provided by research institutes and national resource centres, such as NCBI’s E-Utilities(Sayers et al., 2023) or EMBL-EBI’s Job Dispatcher(Madeira et al., 2024), each again have web UIs and APIs accessible via their own dedicated EaaS clients. Whilst highly capable, Galaxy and other systems provided by national facilities are complex to install and maintain; and are not intended to be deployed to provide access to a few tools needed to enact a workflow.

Conversely, Webias(Daniluk, Wilczyński & Lesyng, 2015) demonstrates how a reusable EaaS platform can provide the same capabilities offered by “bespoke” web sites with all the added advantages of a standardised code execution system. Like Galaxy, it provides an XML based model that describe an executable’s metadata, configurable parameters, and how those parameters are inserted into a command line template. It also offers a simplified grammar for web form creation (buttons, dropdowns, free text entry and file uploads). Results from the program’s execution are presented via a custom web page specified as a Jinja template, allowing “bespoke” views such as interactive JavaScript based visualisations. Whilst Webias’ focus is the easy creation of interactive web interfaces rather than an EaaS web API, it demonstrates that a reusable EaaS can be achieved without complex software frameworks.

Another notable example is cli2rest(Zok, 2026), which was demonstrated with RNAtive, an RNA 3D structure analysis platform(Pielesiak et al., 2025). This EaaS focuses on ease of deployment, by providing an EaaS server designed to be packaged alongside its executable tools within a Docker container. The server’s API allows standardised tool execution via a simple command line client, assisted by a service definition file.

Other systems solely designed for EaaS include OPAL(Krishnan et al., 2006), utilized by Chimera and ChimeraX molecular graphics platforms (Pettersen et al., 2004, 2021), and JABAWS(Troshin, Procter & Barton, 2011; Troshin et al., 2018), primarily employed by users of Jalview (Waterhouse et al., 2009; Procter et al., 2021) to execute a range of alignment and analysis programs. JABAWS has operated at the University of Dundee since 2010, and carried out >80 million jobs/year, and was designed to be easily installed by anyone wishing to take advantage of their own computational resources. However, after over a decade of service, we decided to develop a new EaaS system to replace JABAWS. This new system: Slivka, needed to be easily maintained, and allow any kind of executable command line tool to be provided as a scalable REST web service, without the need to write additional code.

A graphical overview of Slivka is shown in Figure 1. It provides a lightweight “low-code” re-usable framework for creation of scalable, semantically annotated and interoperable web services that allow execution of command line programs. At the back end, server administrators can configure Slivka to run programs locally or according to data size and parameters, schedule them for execution on clusters or other computational platforms. Slivka’s services employ W3C standard HTTP codes and headers, enabling their use by any web client, and MIME types can be specified to maximise interoperability(Rzepa, Murray-Rust & Whitaker, 1998). However, to further simplify integration we also developed Python, Java, and JavaScript client libraries that allow Slivka services to be easily incorporated into other systems. We demonstrate Slivka’s capabilities by creating Slivka-bio, a Slivka service suite for bioinformatics that is provided both as a public server and can be easily installed locally. Slivka-bio services described with EDAM(Ison et al., 2013; Black et al., 2022) annotations allow them to be discovered automatically and accessed by users of the Jalview (Waterhouse et al., 2009) interactive multiple sequence alignment, analysis, and visualisation platform. Slivka was also evaluated in an industry setting, and for this we developed Czekolada – a general purpose Slivka client that runs as a browser application.

**Figure 1.**
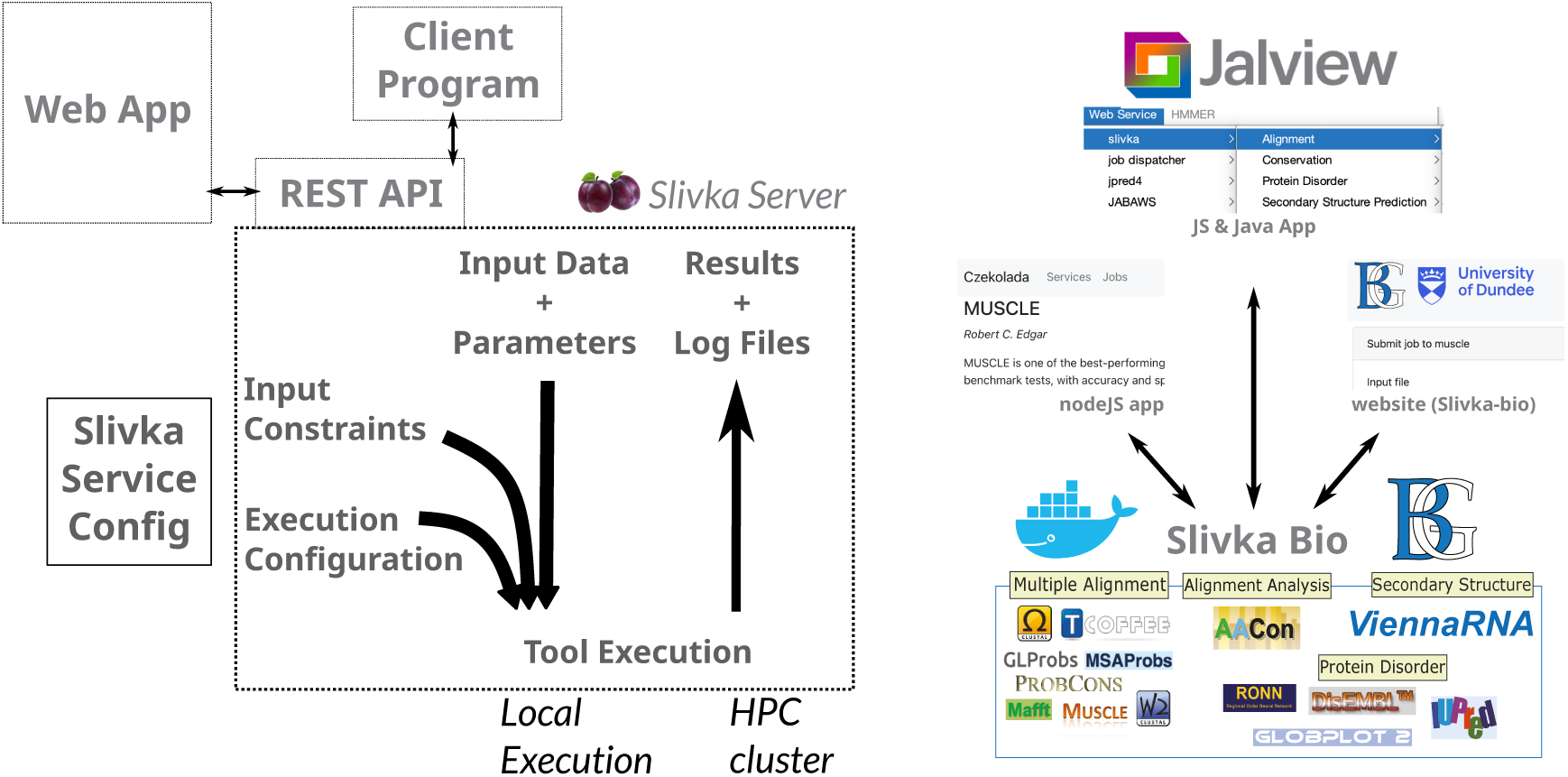
***Overview of Slivka and Slivka-Bio.*** The Slivka system (left) provides a standard REST API that allows remote execution of code for a given set of input data and parameters. Constraints can be defined for a service such as limits on the size of input. and these can also be used to route jobs and tailor runtime parameters for different execution environments. Slivka-bio (right) is a Slivka-based server that provides access to a range of bioinformatics tools, that can be accessed from Jalview and Czekolada, a graphical client for Slivka services. Slivka-bio can be easily run locally through docker, and a public instance is provided by the Barton Group’s server hosted at the University of Dundee. **Figure 1 Alt-Text:** A two-panel figure – the left panel is a schematic representation of a web app and client program interacting with a Slivka server through its REST API. The Slivka server is annotated with the project logo: two purple plums. An additional box shows a service configuration outside the server. Inside the server box three arrows indicate input data and parameters from the web app or client program, input constraints and execution configuration combine to execute a command line tool locally or on a cluster, and another indicating job outputs and logs being returned to the server. The righ panel shows a box labelled with Slivka-Bio, a whale and container docker logo and barton group logo. The box contains several logos and labels: multiple sequence alignment, alignment analysis, protein disorder and secondary structure prediction. Double headed arrows connect it with three labels – nodejs app, JS & Java app, and website (Slivka-bio), each has a screenshot above.

## 2. Design

Our goal with Slivka is to allow easy creation of services that involve execution of command line programs. We also require the system meet the following goals:

- Minimal reliance on specialised web technologies.
- Easy installation and maintenance.
- Low code service configuration.
- Interoperation with existing middleware for workflow and batch execution
- Efficient interchange of data between services
- Allow domain-specific semantic service discovery

Our starting point was to develop a model for EaaS, and a way to express it as a YAML document. This is described in more detail below. The YAML document provides the ‘low code’ framework for creation and configuration of services. It also allows annotation that can be used to convey the function of the service. Slivka provides a ‘discovery’ API that returns a JSON document that lists available services, including their annotations, inputs, and outputs, allowing semantic service discovery. During implementation of Slivka, ‘REpresentation of STate’ principles were employed in the design of its web-API, so services can be called with general HTTP clients. Standard HTTP Status codes are also employed to indicate whether a job has been accepted for processing, and whether results are available for download. Furthermore, uploaded data or results produced by a service are assigned obfuscated identifiers that can be directly used as input to another services. This facilitates efficient data interchange between services and allows callers to enact a complete workflow without download of intermediate files.

### 2.1 A model for the execution of command line programs as services

Slivka’s EaaS model specifies how the execution of a task, such as a command line program or script, can be described as a web service. To do this, the model separates the specification of input data and parameters from details about how the tool is installed and executed. On the server side, the model it allows generation of a series of specific executable commands (essentially a script) tailored to a particular execution environment, for any combination of valid input data and parameters. Callers need only provide data and parameters relevant to the analysis task at hand; any additional parameters or environment set up is provided by the service creator. For instance, system administrators may wish to optimise a service’s execution on high performance (HPC) or virtualized compute platforms through custom parameters and environment variables.

Whilst other EaaS frameworks allow similar functionality, a unique aspect of Slivka (to our knowledge) is that service providers can define and evaluate constraints that may involve programmatic analysis of input data. We demonstrate this in Slivka-bio, by providing functions that allow the number and average length of sequences submitted to an alignment service to be calculated with BioPython(Cock et al., 2009). These serve to validate input and enable a particular environment to be selected according to the size or type of input. The model also includes sections for criteria that allow the EaaS to determine if a particular command line execution was a success, and identify any relevant output files, along with their format. A schematic overview of this process and the associated model components required is shown in Figure 2. See also S1 in Supplementary Materials for an expanded discussion.

**Figure 2.**
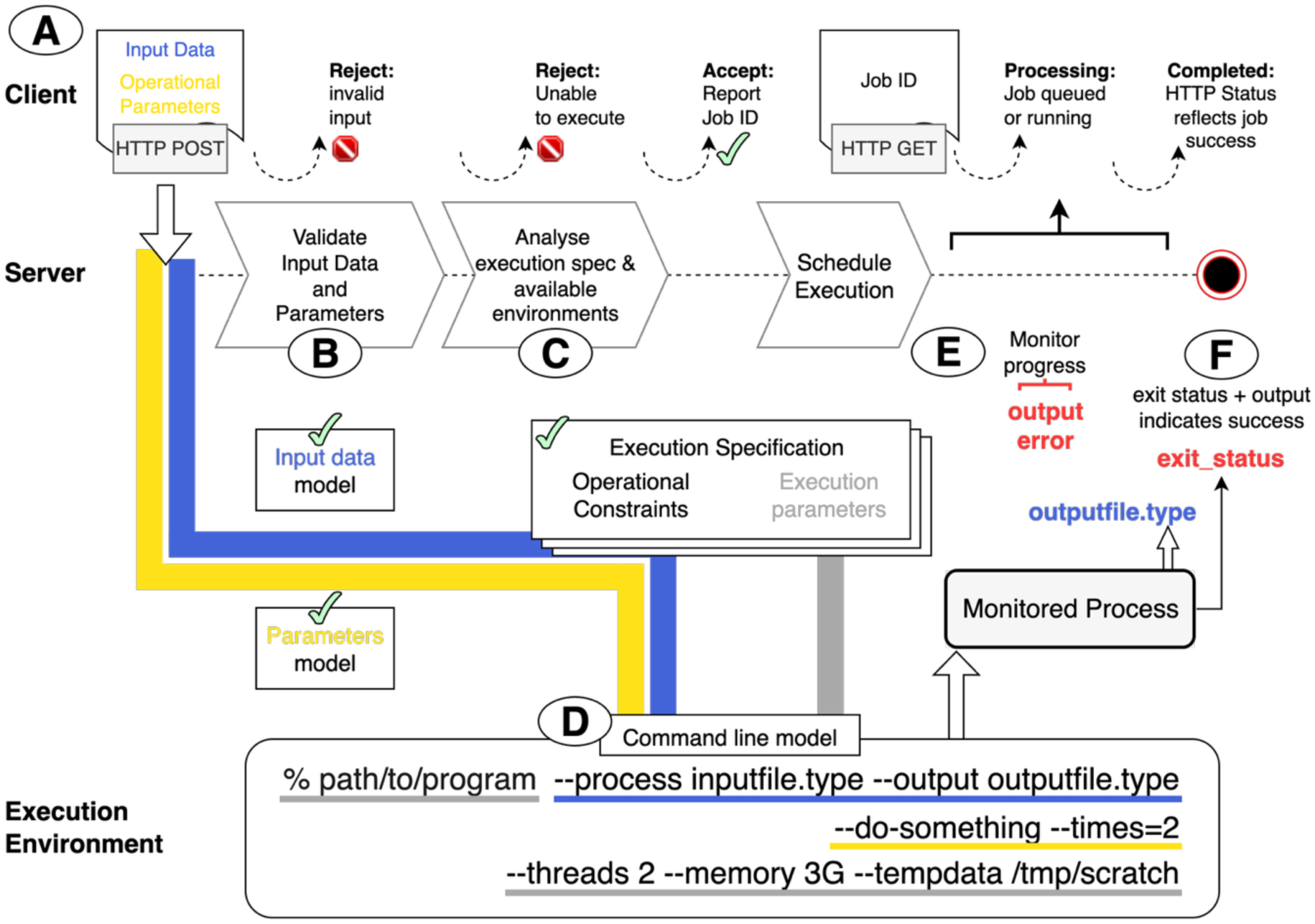
*Slivka’s model for Execution as a Service*. An executable definition consists of models for input data (blue), operational parameters (yellow), expected outputs, a command line model, and one or more execution specifications consisting of operational constraints and environment specific execution parameters (grey). (A) Execution begins with input data (blue) and parameters (yellow), submitted as an HTTP POST request. These are validated using the Service definition’s input data and parameter model (B), and if accepted (tick) matched (C) against the service model’s Execution Specifications. If a specification can be found (tick) where the given input and parameters lie within operational constraints, the system accepts and queues the job. Once one of the eligible executors is available, a tailored command is constructed using the command line model (D). Absolute paths are resolved for input and output files (blue), and arguments are generated for operational (blue) and execution environment specific arguments (grey). (E) Once launched, the process is tracked and status monitored by the client with HTTP GET requests. Standard output or specified log files may also be routed to provide additional progress information. (F) When the process is complete, the system refers to the program specification to determine whether the command line execution was successful, or an error was encountered (e.g. based on POSIX exit_status and generation of a non-empty ‘outputfile.type’). Generated files and logs are also made available for download. See Supplementary Material - S1 for a detailed description. **Figure 2 Alt-Text:** A diagram showing information flow between a client at the top, and the server at the middle, and an environment where data is stored and tool execution takes place at the bottom. Information flows from top left with an HTTP post request that contains input data and parameters which pass through checks in the server based on the services input and parameter data model and execution specification (which includes operational constraints). If all tests pass, they become arguments for a command line in the execution environment, and the client receives a success response. Client HTTP GET requests after this result in status updates, and once the process completes, the result of the tool execution by the server monitoring the process after checking for generated output files and POSIX exit status code.

### 2.2 Semantic service definitions, specification matching, and interoperable systems

Slivka’s YAML service definition files provide the information for ‘service discovery documents’ produced by Slivka’s service discovery API. These documents specify the required input data, their types and any data specific constraints. They also allow inclusion of both human readable free-text and formal annotations, such as those provided by EDAM(Ison et al., 2013; Black et al., 2022). In this way, they serve a second key goal: that the required inputs, outputs, and supported function of services provided by Slivka are sufficiently described to enable specification matching(Zaremski & Wing, 1997), and so facilitate discovery and service composition.

Controlled vocabulary terms such as the EDAM ontology together with typed input and output specifications form implicit contracts. These ‘specification contracts’ allow client systems to automatically select and inject a service according to its function. For instance, many programs for multiple sequence alignment take as input a set of sequences and generate an alignment as output. This pattern of operation is specified by EDAM term ‘operation_0292’ (https://edamontology.github.io/edam-browser/# http://edamontology.org/operation_0292). A client system requiring an alignment service thus only needs to search for services annotated with the corresponding EDAM term to identify those with an input that can take two or more sequences and produces as output an alignment of the input sequences.

### 2.3 Service security and Execution Validation

Security is a principal concern for any system that initiates server-side processes with arbitrary data, and critical when the system is exposed as a public service on the internet. Like any service that acts as a gateway for data exchange with unauthenticated clients, Slivka employs language level Taint Checks (Stein, Lincoln & Stewart, John) to ensure data sent by the client is validated, and prevent unsanctioned code from being executed. Slivka’s EaaS model is also designed to limit exposure of internal data such as authentication keys or logs that might contain sensitive information. Assignment of obfuscated IDs to uploaded files and job results prevents clients from inferring their real location on the server system. Execution limits minimise the impact any client might have on the availability of the service to others; for instance, by exhausting available compute resources. Slivka’s Execution Validator, however, provides the primary mechanism for discarding execution requests that would lead to invalid or otherwise undesirable execution.

Validation must take place at the point of receiving the execution request. If validation fails, human readable error messages are generated. This is an important factor for usability, since by informing the client that input data or parameters lie outside the range of allowed data types and values, the caller can adjust those inputs and submit them as a new job. Execution Validators must therefore be efficient and, ideally, able to run in parallel such that execution requests from multiple clients can be handled in a scalable manner. Slivka’s parameter model allows constraints to be defined and efficiently evaluated for a given input parameter set.

Domain specific functions provided by service developers are critical for verification that uploaded data adheres to the expected format, and we demonstrate this with Slivka-bio’s use of BioPython(Cock et al., 2009) to validate and enumerate sequence data.

### 2.4 Ease of Administration and Server Resilience

Distributed systems are complex to operate and maintain. Servers are routinely restarted, often automatically, in-order to install security patches, update service configurations, or address hardware failures. Slivka implements a decoupled multi-component architecture that provides persistent storage of execution requests and process status that can be read from and written to with minimum blocking. Whilst several industry standard database systems exist that support concurrent access, we opted for a lightweight approach – file system and MongoDB events, in-order to synchronise and coordinate access.

#### 2.4.1 Service resource configuration

Resource allocation and process limits (e.g. memory usage and maximum runtime) are essential aspects of runtime environment configuration. They are, however, highly specific to available infrastructure. We developed an extensible API that allows individual execution platforms to be configured as ‘Runners’. Slivka’s YAML documents can then be used to define general limits for a particular infrastructure, or allow customisation for specific codes, e.g. those that can take advantage of multiprocessors, GPUs or cloud execution systems.

## 3. Implementation

The Slivka execution as a service framework was implemented as a Python 3 package. Python is the most popular language according to the TIOBE Index(“TIOBE Index”), and was chosen due to its maturity and wide use in computational biology. The system has two main components (see Figure 3 and Supplementary Figure S2): the REST Server and Scheduler. They operate on a shared file system and communicate by polling a shared MongoDB instance. The NoSQL database provides persistence for requests, job state, and logical mappings between uploaded data and physical files located on the shared filesystem.

**Figure 3.**
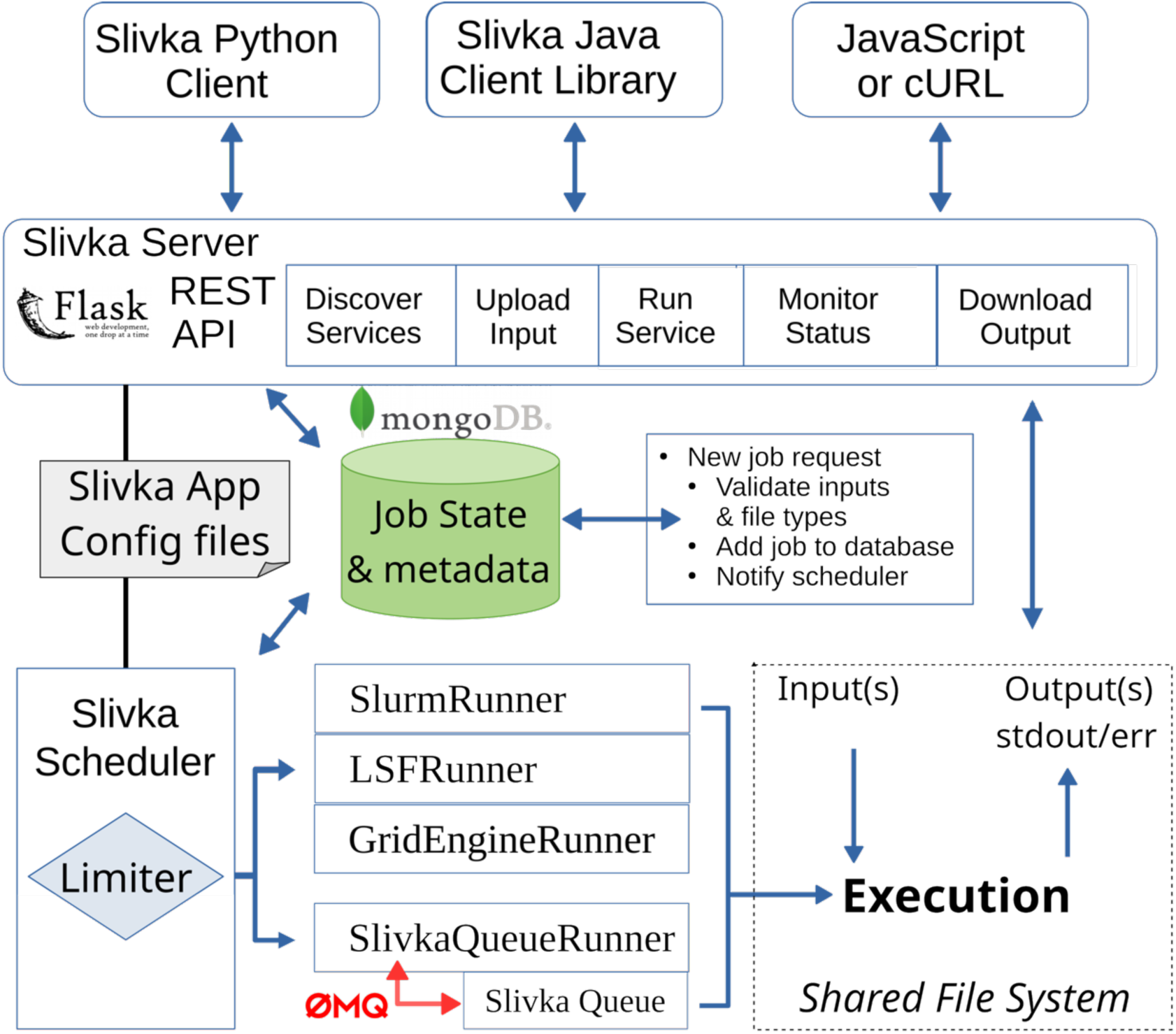
The slivka system. Slivka consists of a Flask based REST server and a Scheduler, linked by configuration files and a shared Mongo Database. Clients for the Server are provided in Python, Java and JavaScript and the server can be accessed directly via HTTP clients such as cURL. API endpoints allow service discovery, data upload, service execution, status monitoring and results download. The server passes valid jobs to the MongoDB which are picked up by Slivka’s scheduler, which passes jobs on to available runners via the limiter, which processes any defined execution environment specific limits. Runners are python classes that implement a standard interface. The local Slivka queue communicates with its Runner via ZeroMQ. Runners for Slurm, LSF and Univa Grid Engine employ command-line tools provided by their respective queuing system to schedule jobs and monitor status. **Figure 3 Alt-Text:** A block diagram running from top to bottom and left to right. Three blocks at the top represent different types of client. Arrows connect the clients with the Slivka server shown in the next row. The server box shows a logo for the flask system and is labelled right to left with boxes for the different parts of the REST API. Next, a rectangular document shape represents Slivka configuration files, and a green cylinder represents a mongo database holding Job state and metadata. Lines and double ended arrows connect these with the rest api, and a double ended arrow lists stages handled when a new job request is received. The next layer shows the scheduler as a box containing a diamond shape representing the limiter logic which is connected by arrows to boxes for each of the job runners, with a red arrow linking SlivkaQueueRunner to the queue process. On the bottom right a box shows the tool execution on the shared file system with arrows shown inputs coming in to the tool execution, and arrow output representing outputs. A double ended arrow connects the execution box with the REST Server layer indicating communication of state and retrieval of job outputs.

### 3.1. Core Architecture

Slivka’s architecture was developed through a series of incremental design-implementation cycles, where different aspects of the system were tested and revised to meet operational needs. We initially considered a heterogeneous distributed architecture unified by a shared SQL database instance, as employed by Webias(Daniluk, Wilczyński & Lesyng, 2015). However, this approach proved unsuitable because it required the database to support concurrent access and update patterns to cope with highly variable response times amongst components in its operational environment. For instance, when a job is selected by a runner for execution, its entry in the shared database should be locked whilst it is submitted to the runtime platform.

Schedulers can sometimes take several seconds to respond to such requests, meanwhile the lock must be maintained, preventing processing of other requests. Although Mysql and Postgres support concurrency, they add additional administration overhead. Messaging busses such as ZeroMQ can also be used to avoid the need for database locks, by allowing time critical state changes to be shared directly between components. However, MongoDB’s support of atomic updates and concurrency obviated the need for additional semaphore mechanisms.

### 3.2 Slivka Applications and the Slivka REST API

Slivka is designed as a generic service provider system and requires at least one service definition to initialise. Service definitions are used to create endpoints for service discovery, data upload, job submission, monitoring and output file download, summarised in Table 1. The Slivka python package provides a command line tool ‘slivka’ that controls slivka’s server and scheduler processes; and allows creation of necessary files and directories based on a predefined template. On start up, the Slivka server and scheduler examine the ‘SLIVKA_HOME’ environment variable to locate a configuration file (settings.yml), and directories for service descriptions (in ‘services’) and application specific python scripts that provide data validators and execution limit functions (in ‘scripts’). The configuration file specifies the locations for uploaded media, and paths where directories for each service execution will be created, along with connection details for the shared MongoDB. Once initialised, the scheduler will immediately start to schedule pending jobs and any periodic test jobs defined for services to verify they are operational.

**Table 1.** Slivka’s HTTP API endpoints.

| URL | HTTP Action and description |
| --- | --- |
| /api | report server version |
| /api/services | Describe all available services |
| /api/services/{serviceName} | describe parameters supported by the service |
| /api/services/{serviceName}/jobs | POST submits a job.<br>202 response on success |
| /api/services/{serviceName}/jobs/{jobid} | GET to retrieve status, DELETE to cancel |
| /api/services/{serviceName}/jobs/{jobid}/log | Retrieves standard output – supports HTTP RANGE queries to allow paged retrieval. |
| /api/services/{serviceName}/jobs/{jobid}/files/{fileid} | URIs for result files |

### 3.3 A Minimal Slivka Application

Slivka’s documentation provided a comprehensive tutorial on the creation of new Slivka services (https://bartongroup.github.io/slivka/getting_started.html#a-minimal-application). For brevity, we only highlight key features of the example service described in the tutorial. Slivka’s reference manual describes these aspects in full (https://github.com/bartongroup/slivka/blob/ver/0.8.5/sphinx-docs/specification.rst).

Executing ‘slivka init new-slivka-app’ creates a new directory containing the Example Slivka application. This consists of an example service definition (example.service.yaml in the ‘services’ sub-directory), the python script that the example service ‘example-service’ executes when it is run, and additional python classes and runner configuration files that are referred to by the service definition’s YAML document. The next few sections highlight aspects of Slivka’s EaaS model as demonstrated by this example service. Supplementary S3 provides the complete service specification including comments and additional description.

#### 3.3.1 Service description file specification

Slivka’s YAML service definition files capture all necessary details of the EaaS definition model introduced earlier Section 2.1, and Supplementary S1. The design was created by analysis of the XML schema components from the JABAWS system, together with the Common Workflow Language (CWL) (Crusoe et al., 2021) tool specification. CWL advocates the use of JSON-LD style macros to import commonly used stanzas, such as installation specific environment configurations, and Slivka’s YAML definitions allow a similar approach. The final version of the schema shown in Figure 4. YAML was chosen not only for clarity and convenience, but also because it allows formal specification with JSON Schema (https://json-schema.org/). When Slivka’s server application is started, discovered service definitions are first validated against the schema to ensure they are well formed. Service parameter, argument and output definitions are however specified as free-text strings in the schema and instead validated with custom parsers. This allows useful feedback to be provided to service developers rather than simply reporting that a service definition failed the schema validation process.

**Figure 4.**
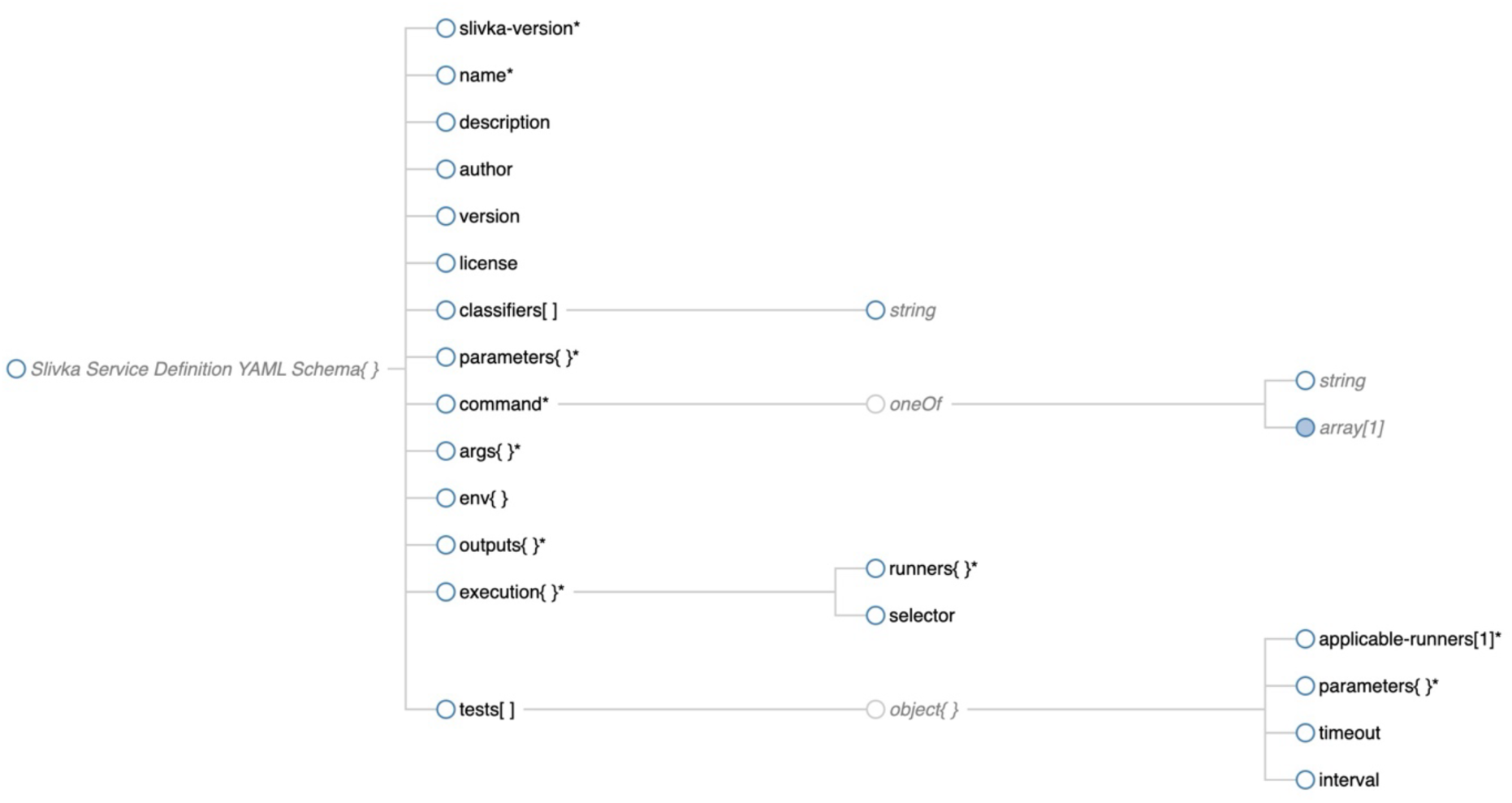
Diagram of Slivka’s Service definition JSON Schema. Service definition files consist of a single root object. Of the 14 sub sections, a definition file must at least contain ‘slivka-version’, ‘name’, ‘parameters’, ‘command’, ‘args’, ‘outputs’, and ‘execution’ (* indicates required sections). Schema diagram generated with json-schema-viewer (Bradley, 2025) **Figure 4 Alt-Text.** A node-linked tree diagram rooted at the left with the base schema node. The next level lists different sub-sections, some of which are marked with a * indicating they are required, and square brackets or curly races indicating they are lists or key:property hashes. The next level shows string type for the classifiers array, a constraint ‘one of’ for command which is either string or an array of strings. A set of named runners and a selector function for the execution section, and for the tests list a set of properties defining applicable runners, the test job parameters, timeout before failing the test, and interval for how frequently to run the test.

The schema for Slivka service definitions is shown in Figure 4, rendered with ‘json-schema-viewer’(Bradley, 2025). The root Slivka object consists of the following elements:

- The version of Slivka that is required to support this service definition.
- metadata describing the specific software executed by the service and any semantic classifiers.
- parameters – which defines input files, flags or other command line parameters provided by callers of the service.
- a command, argument and environment model that defines server-side parameters and flags, and how they are formed into a syntactically valid command line that can be executed by a suitable environment.
- outputs – which allow the specification of the logical name and MIME type of any files or POSIX output and error streams produced during execution.
- execution: which specifies which runners support execution of this service, any environment-specific configuration parameters needed for the runner or program being executed, and constraints (expressed as python functions) that allow environments to be excluded based on provided parameters or size of input data.
- tests – which describes a set of input data and parameters that can be periodically run to ensure the service is operational.

#### 3.3.2 Service metadata

Services are assigned an id based on the filename used for their definition: e.g. example.service.yml generates a service with an id called ‘example’. Definition files must at least contain a name; optional attributes include description, author, version, and license. One or more classifier strings can also be added, for example to allow services to be described using controlled vocabularies (see Figure 5A).

**Figure 5.**
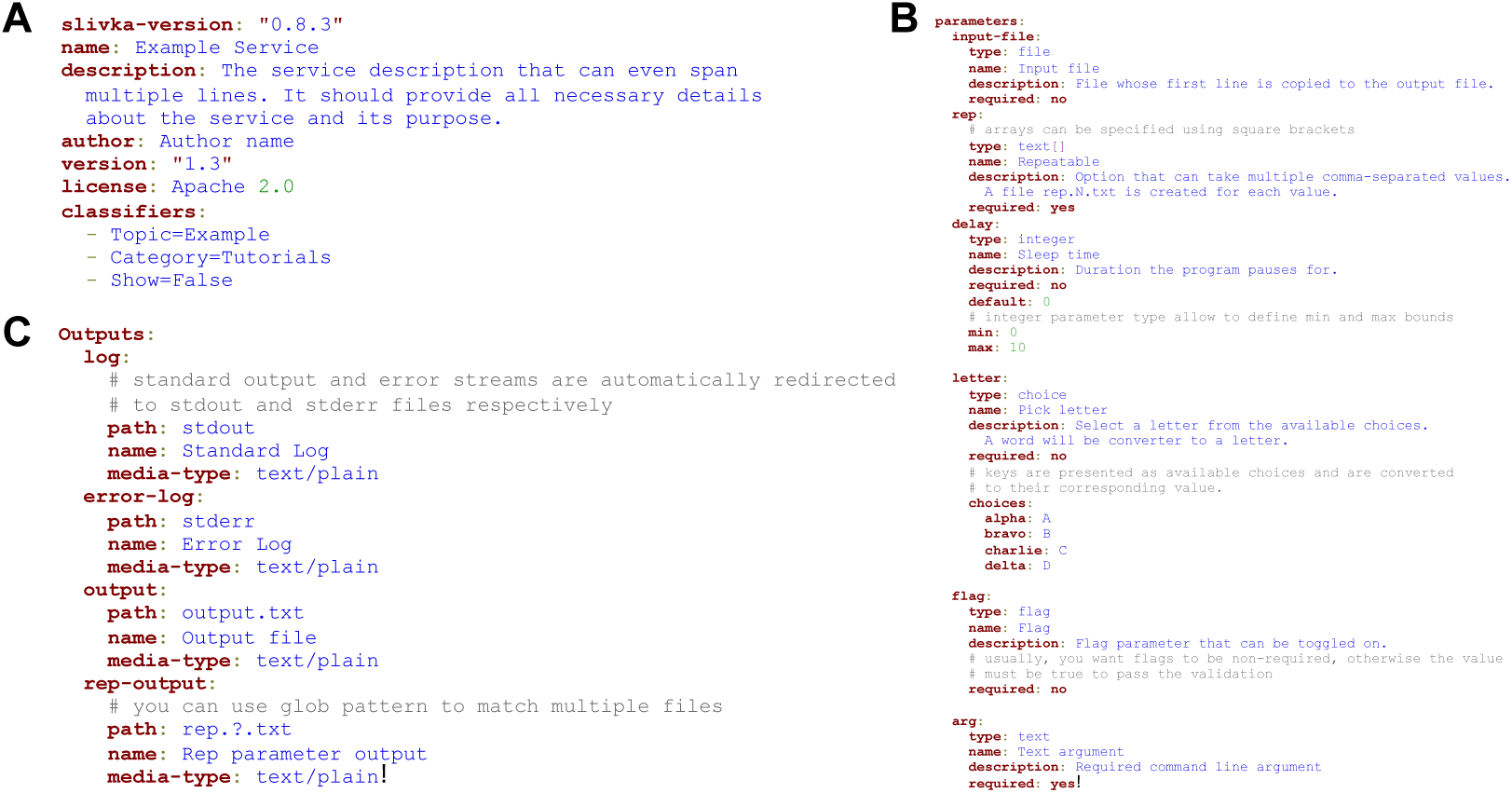
Excerpts from a Slivka service definition file. (**A)** Slivka services allows a human readable name to be assigned, and additional annotations added, including semantic metadata. **(B)** Parameters can be simple values such as plain text or numbers, arrays of values, or files (uploaded or an existing file produced by another service). **(C)** The outputs section allows standard POSIX error and output streams to be exposed to clients, along with any specific files generated during execution. Like input files, outputs are also assigned MIME types so they may be used as input to other services. **Figure 5 Alt-Text:** Three sections of text showing the contents of a service definition file from top left labelled A, B on the right, and C on bottom left.

#### 3.3.3 Input parameters

Input data and adjustable parameters are specified in the parameters section in the YAML service definition file (Figure 5B). Constraint definitions are also possible; for instance, to ensure that mutually exclusive optional parameters are not accepted. In addition to uploaded files, parameters can be specified that take strings, integers, floats, flags, and multiple alternative options. Built in functions allow validation, and these are reused in later sections to map these arguments specified in the service definition to the arguments in the generated command-line. Extensions can also be created to verify uploaded files conform to a specific format or size limit.

#### 3.3.4 Output files

A service’s outputs (Figure 5C) can be discovered and downloaded by clients for a running service. These include standard POSIX output and error streams, and MIME-typed files that can be routed as input to other services for processing.

#### 3.3.5 Command line model

The executable command line is specified via three YAML elements (see Figure 6A).

- ‘env’ – a list of environment variable definitions which can include expressions evaluated according to service input parameters.
- ‘command’ is an array of strings that will be used to generate the command string that will be executed.
- ‘args’ is an ordered list of named properties that defines how inputs and parameters are formed into the final base command string.

**Figure 6.**
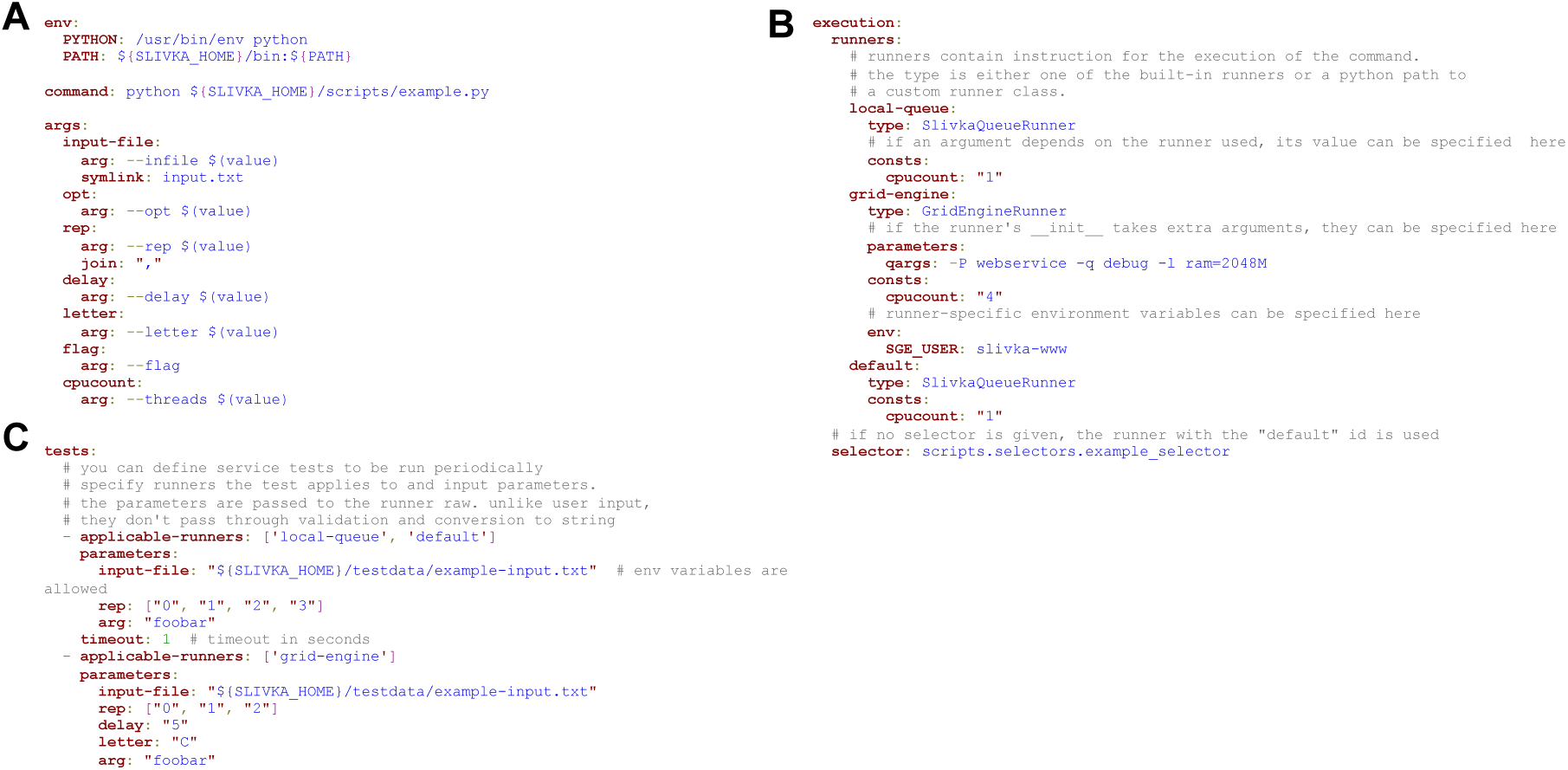
The executable service definition elements. **(A**) Input parameters are mapped with the ‘args’ sub-element to expressions that can be inserted into environment variable definitions (env) and a series of one or more command line strings (command). **(B)** The ‘tests’ section allows sample service inputs to be specified. **(C)** Execution environment configurations can each have their own ‘consts’ definitions, allowing Runner specific generation of commands via the ‘arg’ map. Applicable environments for input data are determined by evaluating a ‘selector’ – a python function found in the scripts directory that is called to determine which environment can support execution of the service for any given input. **Figure 6 Alt-Text:** Three panels of text shown excepts from a YAML service definition file from top left, top right to bottom left.

The ‘args’ list allows incorporation of both input parameters and arguments to be injected that are not specified in the ‘parameters’ section. For instance, in Figure 6A, the ‘—threads’ argument will only be included in the generated command line if the assigned execution environment provides a value for the ‘cpucount’ argument. The args object also allows specification of file name template to accommodate executables that expect a particular ‘dot extension’ (.ext).

#### 3.3.6 generating executable commands and testing executors

Slivka service documents describe how a service’s command line model is translated into a form suitable for execution on one of the available environments. The ‘execution:’ section names one or more runners that can execute the command. These correspond to python classes (see next section). Expressions can be defined for each named runner that are evaluated using the service inputs, allowing injection of data and platform specific configuration details into generated command lines. Execution environments are validated by specification of service tests (figure 6B). Tests for each service can be executed on startup and at specified intervals to ensure execution environments are operational.

### 3.4 Runners, Selectors, and HPC configuration

Slivka Runners are python classes used by the scheduler process to initiate a command line execution and monitor its status. A service’s execution section (see Figure 6C) can specify one or more runners to use for execution by providing ‘selector’ expressions for each runner, along with any additional command line arguments or environment variables needed for execution to the Runner class so it can initiate execution. When the Scheduler picks up a new job from the shared database, the scheduler examines the ‘runners’ section of the service and evaluates these expressions to determine which runner to use. If none are available, the Scheduler will move on to the next unscheduled job. Runners are provided for execution on the local machine, or through Univa Grid Engine, LSF, and Slurm scheduler systems.

#### 3.4.1 Running with containers, workflow and cloud APIs

Runners allow Slivka’s notion of POSIX execution to be decoupled from the underlying service implementation. We developed Runners that allow Slivka clients to use other web service APIs, and to flexibly support container-based execution with minimal change to back-end service configuration. Runners may also reuse other runners, for instance – execution of a container for a tool requires specification of container execution environment, and scheduling of the POSIX commands to manifest the container via one of Slivka’s HPC runners.

### 3.5 Application packaging and dependency management

Packaging and deployment are managed through PIP, which provides a convenient model for management of version-specific dependencies. Domain specific functionality, such as format validators and selectors that provide input data driven functions used in selector expressions requires additional code, also packaged through PIP. Conda-forge’s packaging ecosystem was also be employed to provision command line tools executed by Slivka applications.

## 4. Results

The Slivka framework was created as a domain agnostic EaaS platform that meets the essential and desirable requirements of a secure, publicly accessible web services system (as described in section 2).

### 4.1 Slivka-bio

Our initial goal with Slivka was to evaluate whether it could act as a replacement for JABAWS v2.2, our existing web services framework for bioinformatics that is employed by the Jalview platform. JABAWS consists of an EaaS server, Java library and command line client that together allow a range of open source bioinformatics command line tools to be executed as services.

However, JABAWS employs SOAP style XML services, and their design is tightly coupled to the function of the bioinformatics tools: alignment, alignment analysis, or sequence analysis. There is also no way to run one service on the result of another, because JABAWS’ service API does not allow referencing of results as input to a new execution.

#### 4.1.1 Creation of executable-based services for bioinformatics

The tools provided by JABAWS were looked up in the bio.tools database and, for those not already described, assessed with the EDAM ontology to identify the corresponding operation and expected input and output files. Canonical MIME types for service inputs and outputs were identified, and a set of Slivka validation functions were created to support file content validation with BioPython. Several of the services provided by JABAWS produced output files that did not conform to any standard format. JABAWS implemented custom code to process these files. Instead, scripts were created to transform tool specific outputs into standard formats such as GFF or Jalview annotations files. Rather than create additional data transformation services, these scripts were simply included as part of the execution section in the YAML service definition file.

Table 2 summarises the tools, their inputs and outputs, and the corresponding terms from the EDAM ontology. For tools provided by JABAWS that were not annotated in bio.tools, a minimal set of EDAM terms were assigned. Additional files produced by each program are also made available, along with log files. Custom limit functions for each service implemented with BioPython allow computation of the numbers of sequences and average length. These both validate input files and allow input and parameter dependent executor configuration. Table 3 describes the default limits for each Slivka-bio service, executor limits, and configuration employed for the public Slivka-bio instance at https://www.compbio.dundee.ac.uk/slivka/. The service definition file for T-COFFEE is summarised as a table in Supplementary S4.

**Table 2.** Bioinformatics services provided by the Slivka-bio platform. Each service is listed with expected inputs and outputs, and the key EDAM terms (as full paths from root to term) included in the ‘classifiers’ section of the service YAML file that are detected by Jalview.

| Service | Expected inputs | Expected outputs | Classifiers |
| --- | --- | --- | --- |
| MUSCLE<br>TCoffee<br>MAFFT<br>ClustalO<br>MSAProbs<br>ProbCons<br>ClustalW | Two or more sequences file | Sequence alignment file | Operation :: Analysis :: Sequence analysis :: Sequence alignment<br>:: Multiple sequence alignment |
| GlobPlot,<br>Jronn,<br>DisEMBL<br>IUPred | One or more Sequences | Jalview annotation file,<br>Features file | Operation :: Analysis :: Sequence analysis :: Protein sequence analysis |
| AACon | Sequence alignment file | Jalview annotation file | Operation :: Analysis :: Sequence analysis :: Sequence alignment analysis<br>:: Sequence alignment analysis (conservation) |
| RNAalifold | Sequence alignment file | Jalview annotation file | Operation::Analysis::Structure analysis::Nucleic acid structure analysis::RNA secondary structure analysis::RNA secondary structure prediction<br>Format::Textual format::FASTA-like (text)::FASTA-aln<br>Format::Textual format::ClustalW format<br>Data::Sequence::Nucleic acid sequence::RNA sequence<br>Data::Alignment::Sequence alignment::Nucleic acid sequence alignment<br>Data::Alignment::Sequence alignment::RNA sequence alignment |

**Table 3.**
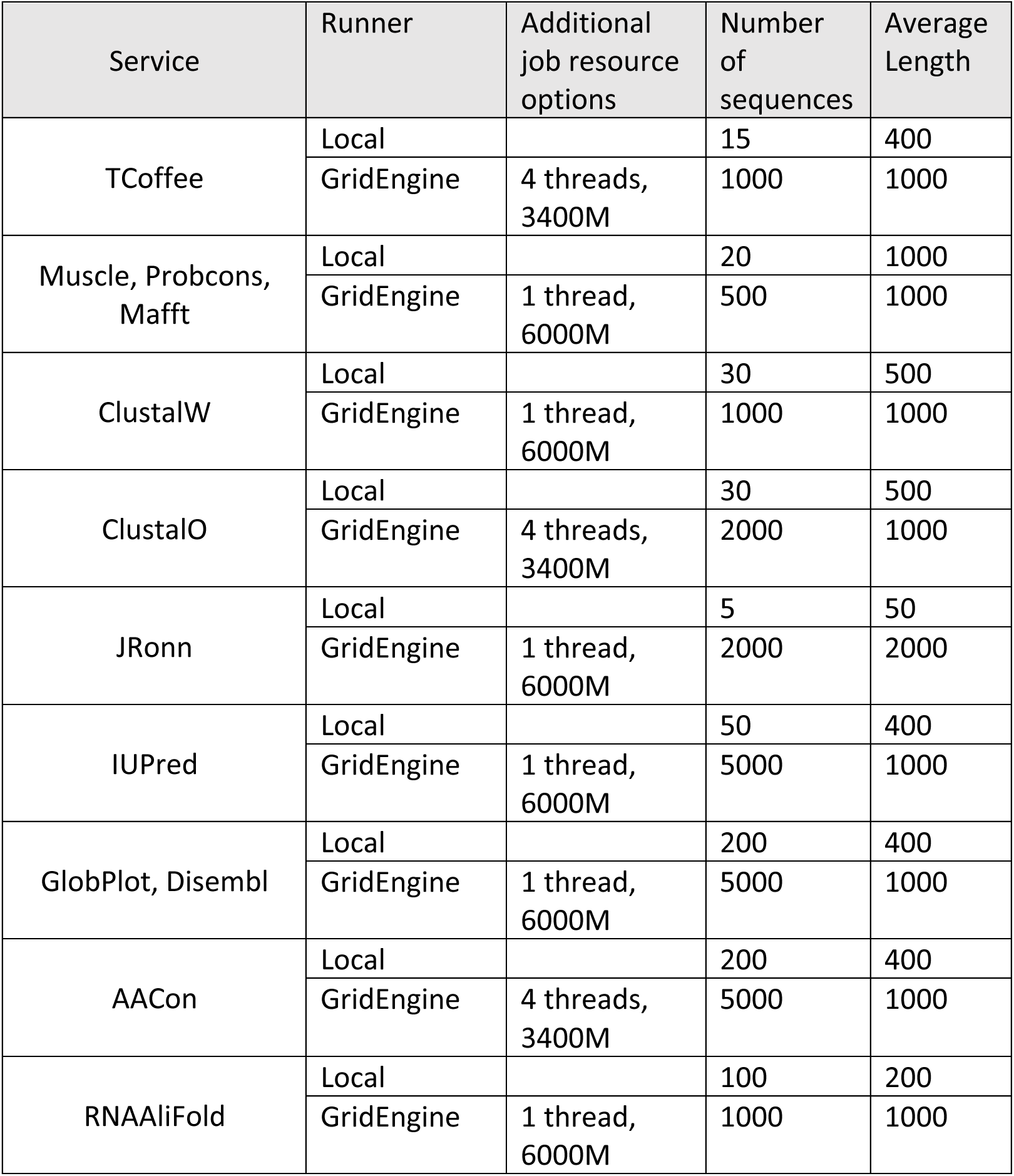
Bioinformatics-specific execution limits for Slivka-bio. For each tool listed in the service column, executable configurations and sequence input limits are given for local and GridEngine based execution for the public Slivka-bio instance hosted by U. of Dundee.

#### 4.1.2 Slivka-bio as a standards-compliant intermediate for third-party bioinformatics services

Web services provided by different systems that perform the same functionality often have very different APIs. For instance, two different web service systems for protein secondary structure prediction may not support the same input and output formats and require different API clients. We explored whether Slivka could enable a single client system to access a web service that normally requires a completely different client: Jpred4, a service provided by the University of Dundee. One approach to achieve this would be to create services that simply wrap an existing command-line client for Jpred4’s REST API. Instead, we developed a Slivka-bio runner that directly interacts with JPred4’s API, avoiding the need to install a Jpred4 command line client and configure an appropriate execution environment. The resulting ‘service specific runner’ demonstrates Slivka can act as a gateway for specialised service APIs, allowing them to be accessed in a unified way.

#### 4.1.3 Distribution and deployment

Our aim with Slivka-bio is to provide a ‘one stop shop’ for anyone wishing to deploy bioinformatics programs as web services. We achieved this by providing two routes to install Slivka-bio: Conda, and Docker, both allowing installation of Slivka-bio and the command-line tools employed by its services.

##### Conda based installation

A Slivka-bio package was created that installs Slivka, and (where possible) all third-party software tools. Most tools in Table 2 were already available through existing condaforge or bioconda channels. New packages were contributed according to community guidelines and licensing, which excluded IUPred(Dosztányi et al., 2005).

##### Docker based installation

A slivka-bio Docker provisioning script was created that 1) installs Slivka-bio from PIP 2) installs all third-party programs via conda and 3) copies a configuration file specifying the location of the Mongo database service and port details for the front end Slivka API. The Slivka-bio system can then be created on any system capable of running x64 architecture code through a docker-compose script which instantiates Slivka-bio and a mongodb container as local services. A docker-compose script was also developed that adds JupyterLab and a ready to use notebook that demonstrates how to discover and access a Slivka-bio multiple sequence alignment service using the Slivka python client.

#### 4.1.4 Service statistics and availability

The public instance of Slivka-bio has been in operation since 2019. The Mongo database instance provides a convenient mechanism for maintaining a statistics database, allowing any Slivka installation to report statistics such as the number and types of services executed over time. When developing JABAWS, we recognised a need to run periodic test jobs to verify all parts of the execution system were operating correctly and disable services that were malfunctioning. A similar functionality was also developed for Slivka, which provides API endpoints reporting which services are available, and execution statistics that exclude any test jobs run during system health-checks.

### 4.2 Clients

Slivka’s HTTP API follows standard HTTP semantics. This means any generic HTTP client, e.g. curl, may be used to discover services, upload input data, and initiate, monitor, and download result files from command line executions. However, additional logic is often needed to parse and process responses, such as service parameter descriptions, when HTTP status codes and header semantics are no longer sufficient. During the development of the slivka-bio application, we thus developed clients in Python, Java and JavaScript that make the discovery and use of slivka services more convenient in these languages. The Python library enables easy operation of slivka services from Jupyter notebooks, whilst the Java library was developed to allow discovery and access to Slivka-bio services from the Jalview platform. Since Slivka is a web-based system, it was also natural to develop a JavaScript platform, Czekolada, that allows job submission, monitoring and results for any Slivka instance through a web-based GUI.

#### 4.2.1 Slivka-client Python library and CLI

The Slivka_client python module is installable via pip from its git repository, or via conda. It provides convenient access to Slivka from other python code or via command line, and employs the ‘request’ library to interact with a Slivka server’s API. The command line interface allows a Slivka server’s services and their parameters to be discovered and reported in human readable form. Jobs can be created by uploading input data and specifying parameters. Once accepted a job’s status can be monitored. Finally, any files and produced from a successful or failed job can be downloaded to a specified directory.

#### 4.2.2 Czekolada

Czekolada is an interactive JavaScript application that provides an easy-to-use web based interface for any Slivka server. Once connected to a server, Czekolada allows users to easily launch and keep track of jobs once they have been submitted, monitor their state, and view or download logs or any generated files. Czekolada is implemented as a React.js application and employs the users’ web browser cache to maintain a record of job IDs for a server. Slivka service definitions are parsed in by Czekolada to generate web forms that allow files to be uploaded and parameters and settings for the service to be adjusted from their defaults.

#### 4.2.3 The Slivka Web Site

Slivka’s web site (www.compbio.dundee.ac.uk/slivka) provides detailed instructions and documentation for Slivka and Slivka-bio. In addition, it also provides web pages that report statistics on the number of jobs run, and the status of the public instance of Slivka-bio that is maintained at U. Dundee. Services can be run directly from the ‘Try it’ pages of the web site: http://www.compbio.dundee.ac.uk/slivka/tryit/, which provides a set of forms dynamically generated from service descriptions. Slivka’s web site is implemented in Python with Jinja2 templates.

#### 4.2.4 Access from Jalview 2.12

Jalview 2.12 provides the most full-featured interface for Slivka services, and integration with jalview was one of the central motivations for development of the framework. A Java client library was first developed to facilitate access to Slivka’s API (https://github.com/bartongroup/slivka-java-client). Jalview’s web services system was then rearchitected to allow services to be accessed via the SwingJS runtime and made available according to their reported EDAM ontology terms.

Jalview 2.12 can query any number of Slivka servers to discover and interact with services annotated with supported semantic terms. Discovered services are accessed through a ‘Web Services’ drop-down menu (shown in Figure 1 below the Jalview logo). This allows Jalview 2.12 users to access Slivka-bio’s services for multiple sequence alignment, conservation analysis, protein disorder predictors and RNA secondary structure prediction. Semantic annotations for services, such as the molecule type for input sequences, allow Jalview to filter the list of services and show only those appropriate for the data being analysed. Constraints associated with EDAM operation annotations, such as the minimum number and length of sequences needed, are also used to filter available services.

### 4.3 Slivka for Drug Discovery

Following development of Slivka-bio, Slivka’s EaaS platform was adopted and evaluated in an industry setting to assess whether it could be used to make specialised tools for structural bioinformatics and drug discovery available to scientists. Computational infrastructure in commercial environments is often different to those in academia, and in order to deploy Slivka, a new runner first had to be developed to allow use of the ‘LSF’ queue system. This new LSF runner code was contributed by Genentech developers to the core Slivka project. Similarly, Czekolada was developed to facilitate use of Slivka services by Genentech researchers. In total, over 80 different tools were published as Slivka services, and more than 1.5M jobs were run.

During this period, we noted that Slivka + Czekolada provide an easy-to-use interface for bench scientists to run computational tools, and aids in recording the provenance of an execution, since all job parameters and code versions are recorded in Slivka’s logs and Mongo database. A minor issue with Slivka’s system was also identified and addressed: initial versions of Slivka tended to create directories containing many millions of subdirectories. Slivka was revised to ensure job storage directories contained a reasonable maximum number of subdirectories at each level to ensure the system remains performant on modern POSIX file systems. Slivka based services for drug discovery are now also accessible through GYDE, a new platform for interactive *in silico* drug discovery data analysis and visualisation(Down et al., 2026).

## 5. Discussion

In our experience, there are two main reasons that researchers provide web interfaces and service APIs for their tools. First, it is both a requirement for publication and an important factor if a tool is to be widely used(Song & Kurgan, 2023). Second, creation of a web application offers an important training opportunity for students and researchers. However, as noted in the introduction, a study of 3649 published services (Ősz et al., 2019) found that on average 15% of services become unavailable 3 years after publication. We tentatively propose that the most common reasons for service attrition are 1) poorly refined service implementations, and 2) lack of personnel and other resources to ensure services continue to operate. Slivka is designed to address the first issue by providing a high quality, hardened and reusable solution for EaaS, and the second, by ensuring the system is easy to administer and maintain without deep technical knowledge.

The secondary goal for Slivka was to create a system that would allow new services to be added for use by clients with a minimum of technological overhead or need to update client code. We demonstrate this with Slivka-bio and Jalview 2.12 through incorporation of EDAM Ontology terms and MIME-typed inputs and outputs. Rather than implement formal standards for interoperability (e.g. WS-I), we adopted a pragmatic, self-documenting approach compatible with standard service catalogues (e.g. bio.tools), drawing requirements and inspiration from our own JABAWS, as well as Galaxy, Nextflow, and The Common Workflow Language. Whilst knowledge of specialised semantic representations is not required to discover and use compatible services, Slivka’s semantic annotation model does rely on *a priori* knowledge. For the case of Slivka-bio, clients need knowledge of how the EDAM operation and data ontologies they are interested in map to the data formats and MIME types of service inputs and outputs. This approach, implemented in Jalview 2.12, enables *specification matching* between services and required inputs, outputs, and operations requested by the client to be accomplished with minimal complexity. At the implementation level, it also allows specifics of an executable providing the function to be hidden behind a standardised interface, further promoting reuse. As a result, Slivka servers support creation of services that enact functional workflows. Data produced from any services can be passed as input to other services providing the types of input and output match, simply by referencing the data’s output URI, and without intermediate download.

Figure 7 summarises the currently available resources in the Slivka Ecosystem. Slivka’s capabilities were initially demonstrated through the creation of Slivka-bio and its integration into the Jalview platform. The Slivka website at University of Dundee provides a web user interface, documentation and up to date status information for the public-facing Slivka-bio installation accessible by Jalview 2.12. Users of the native Jalview 2.12 desktop and JalviewJS web applications may also download and install Slivka-bio for use on their own machines via conda and Docker. Slivka was also employed in the development of LIGYSIS(Utgés, MacGowan & Barton, 2025) and allowed server side computation to be abstracted from web development, enabling the main author to focus on user interface and results presentation (Utges, J. *priv. Comm*). However, Slivka’s adoption at Genentech provided independent verification, and underscored its ease of deployment and readiness for use as a production system. Genentech’s deployment of Slivka allowed biologists access to state-of-the-art computational structural biology tools, initially, though Czekolada’s general-purpose graphical interface, and later with GYDE(Down et al., 2026), an interactive visual analytics platform for drug discovery. It also demonstrated how a unified EaaS platform can facilitate reproducible computational science in an industry setting through provision of a centralised log of tools, input data and results. We also engage with tool developers to create Slivka service definition files for command line tools to facilitate their use via Slivka (e.g. ESMRank(Arnese & Gambardella, 2026)).

**Figure 7.**
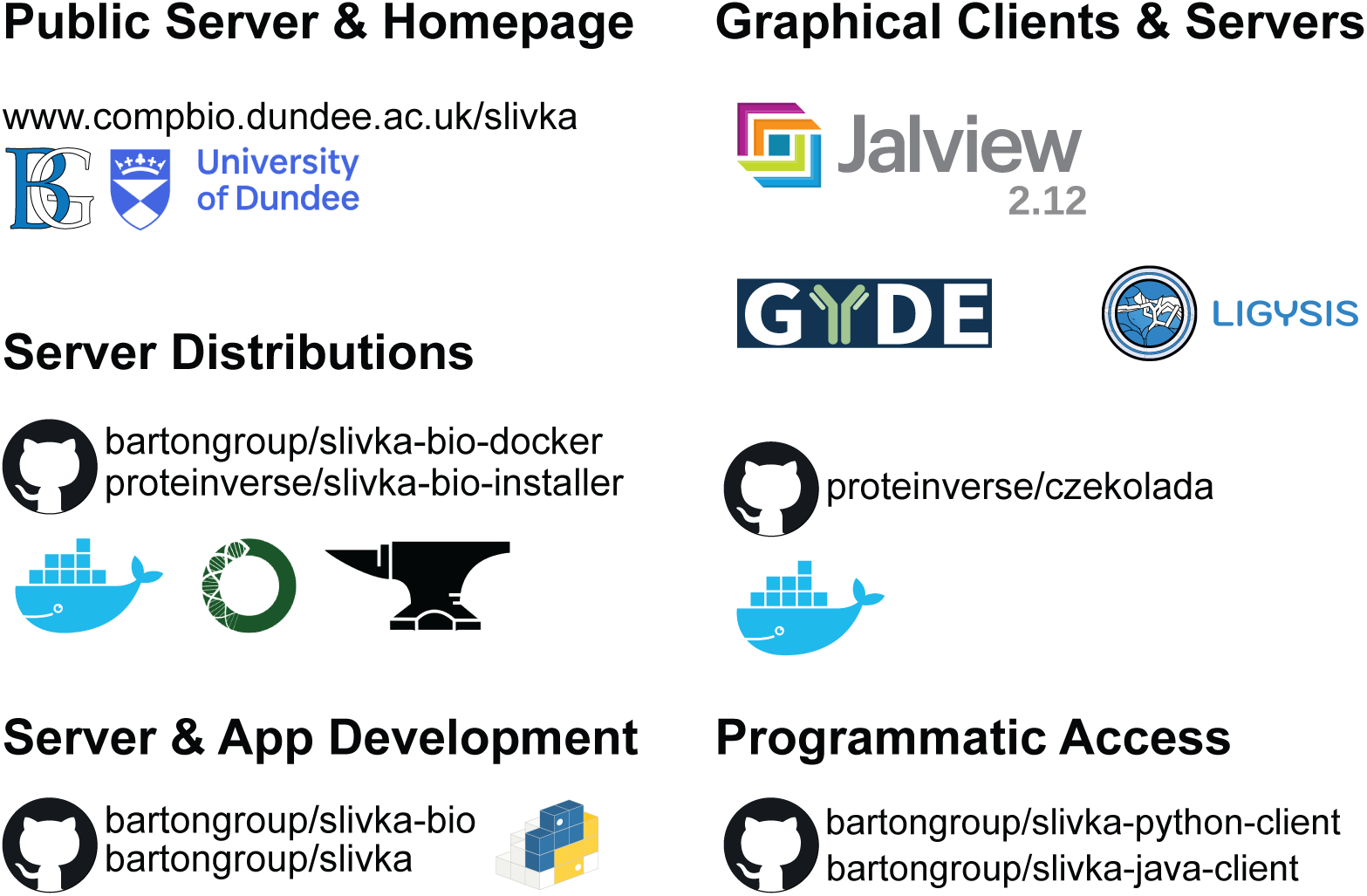
The Slivka Ecosystem. The core Slivka application framework is made available via Conda and PIP, and Slivka clients have been created in python, Java and JavaScript (see czeckokada). Slivka-bio, available via Conda and Docker, provides a ready-to-use Slivka application that allows access to a number of specialised bioinformatics programs, and a publicly accessible instance is available via the Barton group’s homepage, maintained by the Dundee’s Resources for Protein Sequence Analysis and Structure Prediction (DRSASP). DRSASP’s server also has a simple web-based user interface that allows services to be executed and their status monitored. A java client for Slivka has been created and integrated it into the Jalview platform for interactive multiple alignment and analysis as a stand-alone desktop application, command line tool or run directly in the browser as JalviewJS. We also developed Czekolada, a ReactJS based node.js app that allows services to be discovered and used to analyse data. **Figure 7 Alt-Text:** A figure showing 5 sets of items grouped by name, with server addresses, git repositories, and logos from github, docker, bioconda and condaforge, ligysis, gyde, the university of Dundee, the Barton Group and Jalview.

Slivka’s development took place before advent of Agentic AI. However, its HTTP API allows many of the same capabilities offered by MCP, which was introduced by Anthropic in 2024 to allow AI systems to call programs and other computer-based tools. Unlike MCP, however, executables exposed as Slivka services can be accessed with a basic HTTP client, and clients only need to implement logic sufficient to poll for and download program outputs. Such logic can also be described to an Agentic system via Skills(“Agent Skills Specification”), introduced by Anthropic in 2025 that are both more token-efficient and easier to create than MCP. We consider Slivka’s EaaS API to be compatible with both approaches, since its service descriptions employ natural language descriptions and ontology terms for description of operations, and allow MIME types to be specified for inputs and outputs. LLMs are also well suited for the generation of Slivka service descriptions, and we welcome collaborations focused on evaluating how LLMs can aid the generation and consumption of Slivka services.

### 5.1 Conclusions

In this work, we have sought a common language, or medium, that both computers and humans are able to understand, for the execution of computational data analysis tools. Our solution, Slivka, defines an interface that allows programmatic execution of those tools, and together with its various clients, the Slivka API allows humans access to those tools through the same medium. We have demonstrated this through creation of Slivka-bio, that provides services for a widely used stand-alone application, Jalview, and a new web-based interactive platform, GYDE. Slivka is a general system, and not limited in application to biology, so has potential to enable interoperation across very different areas of science. Furthermore, it has already enabled advances in both Academia and Industry, and an open roadmap for future extensions to Slivka’s ecosystem is planned. We invite all who share our vision of “easy to create, and easy to access” services for computational code to contribute extensions and new service definitions via Slivka and Slivka-bio’s github repositories. For the latest updates on Slivka based applications, service additions and other resources, please see https://www.compbio.dundee.ac.uk/slivka.

## Author Contributions

G.J.B. and J.B.P. initiated the Slivka project and secured funding. M.W. designed and implemented Slivka, based on conceptual models from J.B.P., and created the Jalview v2.12 prototype able to access Slivka services from the native application and browser based JalviewJS. K.M. and T.D. developed the application of Slivka in drug discovery. T.D. designed and developed Czekolada, and Slivka runners for Slurm and PBS. S.G. managed integration of Slivka services with the Dundee Resource for Sequence Analysis and Structure Prediction, developed Slivka-bio docker containers, and example jupyter notebooks. J.B.P. managed the project, supervised M.W., and drafted the manuscript.

## Acknowledgements

Robert Hanson (https://orcid.org/0000-0001-5411-2356), developer of SwingJS, provided support to M.W. during integration of Slivka-bio services with JalviewJS 2.12. Fabio Madeira (https://orcid.org/0000-0001-8728-9449), lead developer of EMBL-EBI web services, provided advice and support integrating clients for EMBL-EBI’s HMMER services with Jalview. Help and advice was also provided by the HMMER developer community. Thomas van Aalten originally developed Jalview’s integration with HMMER as part of a University of Dundee Summer Scholarship. We also acknowledge Dr James Abbott (https://orcid.org/0000-0001-7701-4249), co-lead of The Dundee Resource for Sequence Analysis and Structure Prediction (DRSASP), who provided technical guidance to S.G., and the University of Dundee Research Computing team.

## Funding information

This work was supported by The Wellcome Trust [218259Z/19/Z to G.J.B. & J.B.P., 101651/Z/13/Z to G.J.B]; and the UKRI Biotechnology and Biological Research Council (BBSRC) [BB/L020742/1 to G.J.B, BB/X018628/1 to G.J.B, and BB/R014752/1 to G.J.B).

## Notes

### Competing Interest Statement

The authors have declared no competing interest.

