## Supplementary Information for "Slivka and Slivka-bio: a lightweight framework for presenting executables as web services and its application in bioinformatics"

This supplementary information document consists of the following sections:

- S1 – Essential descriptors for command line execution.
    - A semi-formal analysis of the declarations and process functions involved in generation of a command line tailored for execution on a particular POSIX compliant platform.
- S2 - Supplementary Figure – detailed diagram of the Slivka system architecture.
- S3 – Example Slivka Application.
    - The service definition for 'example.service.yaml' – the example service created when a new Slivka server is initialised.
- S4 – Slivka service

### Supplementary S1. Essential descriptors for command line execution

Most operating systems in use today provide a POSIX-like environment that supports command line execution. More specifically, they allow code, packaged as programs, to be invoked by forming a command line. This is simply an ordered series of expressions interpreted by the command line environment that ultimately resolve to a program's location and its means of execution.

Command Line := {<program location & execution spec>, {arguments} }
Executor( Command line ) -> Process
Process := { process_identifier, input stream, output stream, error stream }

An invoked instance of a program is a process – which can communicate via standard input, output and error streams allocated by the operating system. Output streams are typically used to report progress during the program's execution. Command lines also allow additional parameters to be passed to programs, through command line arguments. How these additional parameters are ordered and interpreted is program dependent; but usually parameters include file locations for data import or result export, or allow adjustment of the way the program operates, such as modification of a threshold.

arguments := { *program_ordering* ( {input data spec}, {output data spec}, {operational parameters}, { environment parameters} ) }

In addition to input data and operational parameters, several other factors can affect a program's execution. These are controlled through custom installation and runtime resource settings, perhaps specified as environment variables or system-specific parameters. Such details are for the large part uninteresting to the user of a program, but of great interest to the executing system's administrator, since adjustment may be necessary to take advantage of special hardware, or when handling large input data sets. For instance, many simulation codes allow tuning to specify the use of additional compute or memory resources. Similarly, it is useful to recognise and prevent command line executions where specified parameters or input data are not valid. Whilst programs usually perform such a validation step, not every program is well designed, and for those that are, they may not do so in a timely or compute aware manner.

Execution Validator (..) := { < program input data spec >, < program parameters spec >, <available program execution environments> }
Execution Validator ( { input data }, {operational parameters} ) -> invalid_execution | { customised environmental parameters }

Given a domain specific language that allows the definition of a schema to describe the input data and parameters that a program allows, it is then possible to define a validator function that can efficiently decide whether 1) a given set of input data and parameters are valid for a program, and 2) whether any additional environmental parameters are needed for command line execution.

#### Runtime platforms

One of the most important benefits of EaaS systems is that they decouple runtime specifics from program execution. These specifics are two-fold. Firstly, they include the necessary environmental parameters required for the runtime to execute an installed program with a particular set of input data and control parameters. Secondly, they also include environmental parameters specific to the platform where the executable's process is launched. Different runtime platforms will usually require different environmental configuration to run an installed executable, and some platforms, for whatever reason, may not be able to support execution with a particular combination of input data and parameters. These natures can be encapsulated defining re-usable 'Runners', and for each service, a 'Selector'.

Runner := { identifier, methods for starting, stopping and monitoring state of process }

Selector ( .. ) := { {service}, {input data},{operational parameters} } -> { null | { Runner } }

A Runner represents logic and platform specific configuration data necessary for initiation and management of processes for a command line generated by the Executor. The simplest Runner passes command lines to the local system and allows any resultant processes to be monitored and killed. Other runners may utilise distributed schedulers, or even pass command lines to other systems for execution. Regardless, a Selector is required to map the given set of input data and operational parameters for a service to a set of Runners able to undertake execution.

#### Process status, exit codes and results discovery

Once a process has been successfully created by a runner, its status can be monitored, until it terminates. Under POSIX, the resultant exit status conventionally indicates whether errors were raised, or whether its execution was a success. There may additionally be information from Runners concerning the reason for termination. For instance, because the process exceeded allocated resource limits such as Memory or allowed time. Notwithstanding abnormal termination due to system limits or user requests, once a process has completed, executables can be expected to generate results files in addition to standard output and error streams. Dependent on the nature of the executable, these files may be written to hard-coded paths or according to the {output spec} passed via the command line arguments.

Executor(Runner, {service},{input data}, {operational parameters }) -> Process_Execution

Process_Execution := { status, out_stream, error_stream, exit_code, generated files }

For most executables, the type of the files generated during execution are known in advance, and it is usually safe to assume that if the contents of those files conform to the expected type, the execution was successful. There may also be additional files produced during execution which may not be needed to be returned following a successful execution. To select the results files of interest, it is often most convenient define a filename glob for each result file expected to be produced, enabling them to be recognised and associated with their expected type.

Result_file := { file_name, file_format }
Output data spec := { {Result_file, internal_file_glob } … }
Result_set(..) := Process_Execution -> { exit_code, out_stream, error_stream, {result_file} }

### Content types for input and output files

Most established domains in computational science have tools that produce files in standardised formats that facilitate interoperation with other software used in the domain. MIME types provide a convenient, extensible model for the formal naming of these formats, allowing the content of files provided to and generated by an executable to be unambiguously described.

Whilst a general framework should support the advertising of input and output data formats, it cannot provide implementations of content validation, only hooks that allow validation to be used to ensure correctness. In our experience, we found dynamic validation to be impractical, however, and the implementation we developed does not carry out validation of output data. Instead, our system only allows domain specific format validation for input data, which is the necessary first step to extract domain-specific quantities needed to allow selection of Runners based on provided input.

### A minimal description for an executable as a service

Having defined the component transformations and operations necessary to formally construct, execute, and select results of a command line program, it only remains to describe the specifics for a particular executable so it can be executed given input data and operational parameters. These are:

- Program location & execution spec
    - Template for command line passed to runner
        - Command line arguments
        - Shell Environment settings
- Input data spec
    - Type
    - Validation expression
- Output data spec
- Operational parameter spec
    - Type (e.g. String, integer, float)
    - Validation expression (e.g. minimum/max values for numbers)
- A map for each execution environment between
    - Boundary conditions for allowed input data and operational parameters
    - Runner specific environmental parameters

### Supplementary Figure S2. Slivka Execution as a Service System software architecture.

Figure S2 shows a detailed schematic of Slivka's architecture. The Slivka server process (left) implements the front-end HTTP API, validating data and input parameters for services, saving and retrieving jobs from MongoDB, storing input data files uploaded as input to services and serving files that have been produced during job execution. The Slivka Scheduler (right) manages execution of submitted jobs on Runners for eligible execution environments, polls execution environments to update job statuses in MongoDB, and runs automated tests to verify services are operational. Configuration, selector scripts, MongoDB database and shared filesystem are repeated for clarity.

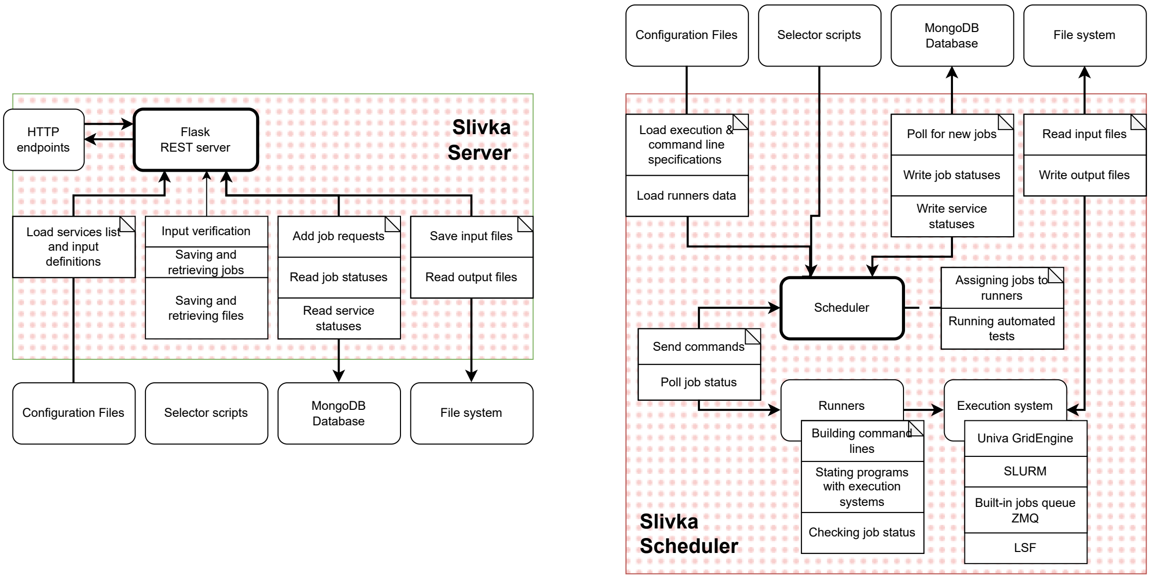


S2 Figure Alt-text: Two software logical architecture diagrams with lines and arrows connecting external components. Within each diagram are square boxes indicating distinct operations with arrows connecting them to the two main processes represented as bold rounded edge boxes.

### Supplementary S3. The Example Slivka Application

Slivka includes an example Slivka application template that demonstrates most aspects of its Executable as a Service (EaaS) model. To create the example application, install Slivka via pip or conda, then execute the command ‘slivka init example-application’ to create a new Slivka application directory called ‘example-application’.

The full version of the services/example.service.yaml file is reproduced below with highlights generated with https://tohtml.com/yaml/ using the ‘rare:YAML’ and ‘White’ settings.

```yaml
# This is an example service configuration.
# Services are specified in the <name>.service.yaml files
# which are automatically detected by slivka when placed
# in the services directory.

---
slivka-version: "0.8.3"
name: Example Service
description: The service description that can even span
  multiple lines. It should provide all necessary details
  about the service and its purpose.
author: Author name
version: "1.3"
license: Apache 2.0
classifiers:
  - Topic=Example
  - Category=Tutorials
  - Show=False

parameters:
  input-file:
    type: file
    name: Input file
    description: File whose first line is copied to the output file.
    required: no

  opt:
    type: text
    name: Text
    description: Some optional text.
    required: no
    # different parameter types have unique properties such as
    # max-length limit for text type
    max-length: 24

  rep:
    # arrays can be specified using square brackets
    type: text[]
    name: Repeatable
    description: Option that can take multiple comma-separated values.
      A file rep.N.txt is created for each value.
    required: yes

  delay:
    type: integer
    name: Sleep time
```

```yaml
    description: Duration the program pauses for.
    required: no
    default: 0
    # integer parameter type allow to define min and max bounds
    min: 0
    max: 10

  letter:
    type: choice
    name: Pick letter
    description: Select a letter from the available choices.
      A word will be converter to a letter.
    required: no
    # keys are presented as available choices and are converted
    # to their corresponding value.
    choices:
      alpha: A
      bravo: B
      charlie: C
      delta: D

  flag:
    type: flag
    name: Flag
    description: Flag parameter that can be toggled on.
    # usually, you want flags to be non-required, otherwise the value
    # must be true to pass the validation
    required: no

  arg:
    type: text
    name: Text argument
    description: Required command line argument
    required: yes

# run the script with python; environment variables are supported
command: python ${SLIVKA_HOME}/scripts/example.py

args:
  # create a symlink and pass its name instead of the real path
  input-file:
    arg: --infile $(value)
    symlink: input.txt

  opt:
    arg: --opt $(value)

  # join provided values with comma
  rep:
    arg: --rep $(value)
    join: ","

  delay:
    arg: --delay $(value)

  letter:
    arg: --letter $(value)

  # the value is either "true" or None and shouldn't be included in the
argument
  flag:
```

```yaml
    arg: --flag

  # this argument does not map to any input, but is populated from the
  # runner defined constants
  cpucount:
    arg: --threads $(value)

  # to add a fixed argument which is not present in the inputs
  # give it a dummy default value, so that it's not skipped
  _separator:
    arg: --
    default: present

  # use the value as an argument directly
  arg:
    arg: $(value)

env:
  # customize environment variables
  # only variables defined here plus PATH and SLIVKA_HOME are available
  # during command execution
  PYTHON: /usr/bin/env python
  PATH: ${SLIVKA_HOME}/bin:${PATH}

outputs:
  log:
    # standard output and error streams are automatically redirected
    # to stdout and stderr files respectively
    path: stdout
    name: Standard Log
    media-type: text/plain
  error-log:
    path: stderr
    name: Error Log
    media-type: text/plain
  output:
    path: output.txt
    name: Output file
    media-type: text/plain
  rep-output:
    # you can use glob pattern to match multiple files
    path: rep.?.txt
    name: Rep parameter output
    media-type: text/plain

execution:
  runners:
    # runners contain instruction for the execution of the command.
    # the type is either one of the built-in runners or a python path to
    # a custom runner class.
    local-queue:
      type: SlivkaQueueRunner
      # if an argument depends on the runner used, its value can be
specified  here
      consts:
        cpucount: "1"
    grid-engine:
      type: GridEngineRunner
      # if the runner's __init__ takes extra arguments, they can be
specified here
      parameters:
```

```yaml
        qargs: -P webservice -q debug -l ram=2048M
      consts:
        cpucount: "4"
      # runner-specific environment variables can be specified here
      env:
        SGE_USER: slivka-www
    default:
      type: SlivkaQueueRunner
      consts:
        cpucount: "1"
  # if no selector is given, the runner with the "default" id is used
  selector: scripts.selectors.example_selector

tests:
  # you can define service tests to be run periodically
  # specify runners the test applies to and input parameters.
  # the parameters are passed to the runner raw. unlike user input,
  # they don't pass through validation and conversion to string
  - applicable-runners: ['local-queue', 'default']
    parameters:
      input-file: "${SLIVKA_HOME}/testdata/example-input.txt"  # env
variables are allowed
      rep: ["0", "1", "2", "3"]
      arg: "foobar"
    timeout: 1  # timeout in seconds
  - applicable-runners: ['grid-engine']
    parameters:
      intput-file: "${SLIVKA_HOME}/testdata/example-input.txt"
      rep: ["0", "1", "2"]
      delay: "5"
      letter: "C"
      arg: "foobar"
...
```

### Supplementary S4. Slivka Service definition for T-Coffee

The tables below are derived from the Slivka-bio service definition for T-Coffee.
https://github.com/bartongroup/slivka-bio/blob/8107ff2de67341b288240c1211d22391a5f8f9fa/services/tcoffee.service.yaml

**Metadata**

**slivka-version:** 0.8.3
**name:** TCoffee
**description:** T-Coffee (Tree-based Consistency Objective Function for Alignment Evaluation) is a multiple sequence alignment software using a progressive approach. It generates a library of pairwise alignments to guide the multiple sequence alignment.
**author:** Cedric Notredame
**version:** '13.41.0'
**license:** GNU GPL
**classifiers:**
- 'Topic :: Computational biology :: Sequence analysis'
- 'Operation :: Analysis :: Sequence analysis :: Sequence alignment :: Multiple sequence alignment'

parameters:

```
  sequence:
    name: input sequence file
    type: file
    required: true
    media-type: application/fasta

  mode:
    name: Preset Mode
    description: '...'
    type: choice
    required: false
    choices:
      quickaln: quickaln
    default: quickaln

  tg-mode:
    name: Terminal gaps penalty
    description: '... '
    type: choice
    required: false
    choices:
      '0': '0'
      '1': '1'
      '2': '2'
    default: '1'
```

```
  iterate:
    name: Number of iterations
    description: ...
    type: int
    required: false
    min: -1
    max: 100
    default: 0

  outorder:
    name: Output order
    description: ...
    type: choice
    required: false
    choices:
      input: input
      aligned: aligned
    default: input
```

outputs:

```
  alignment:
    path: '*.clustalw'
    media-type: application/clustal
  log:
    path: stdout
    media-type: text/plain
  error-log:
    path: stderr
    media-type: text/plain
```

Command & Argument model

```
command:
- t_coffee

args:
  _output:
    arg: -output=$(value)
    default: clustalw
  sequence:
    arg: -seq=$(value)
  mode:
```

Environment, Runners and tests.

```
env:
  TEMP: /tmp/

execution:
  runners:
    default:
      type: SlivkaQueueRunner

tests:
- applicable-runners: ["default"]
```

| | |
|---|---|
| arg: -mode=$(value)<br>tg-mode:<br> arg: -tg_mode=$(value)<br>iterate:<br> arg: -iterate=$(value)<br>outorder:<br> arg: -outorder=$(value) | parameters:<br> sequence: ${SLIVKA_HOME}/testdata/uniref50.fa<br> timeout: 5<br>... |
